# BARCS: beta-binomial regression for multivariable CRISPR screen designs

**DOI:** 10.64898/2026.08.31.748412

**Authors:** Kyu-Won Lee, Hyun-Hwan Jeong

## Abstract

Pooled CRISPR screens increasingly use longitudinal, donor-adjusted, and factorial designs, but beta-binomial screen methods have largely remained limited to pairwise comparisons. BARCS extends the library-total-conditional beta-binomial model to guide-level regression with an arbitrary design matrix, enabling direct estimation of time, covariate, and interaction effects. In four replicate-complete Cas13 screens, adding the intermediate time point modestly improved essential-gene recovery. Applying the same non-targeting-control scaling rule to BARCS, MAGeCK-MLE, edgeR-QL, DESeq2, and limma–voom produced similar calibration across all five methods, while the four alternatives ranked essential genes more strongly than BARCS. In an ordered-bin IL2RA screen, donor-adjusted BARCS recovered more validated regulators with fewer total calls than the matched four-bin MAGeCK-MLE fit, and cross-fitted controls exposed excess guide-level significance. Simulations showed gains from dispersion moderation and control-based denominators, but seed-specific results exposed denominator sensitivity and a null grid localized substantial gene-level error to correlated-guide aggregation rather than dispersion alone. Aggregation-matched control scaling reduced but did not eliminate this error. An external audit prompted by concerns about beta-binomial false discoveries showed that the reported CB^2^ null-discovery count disappeared when full-library totals were restored. This corrected one denominator-dependent result but did not refute the broader calibration concern; nominal-level calibration remained unresolved. BARCS therefore contributes a multivariable extension of the library-total-conditional beta-binomial model together with an explicit account of where its inference is valid: guide-level coefficients are supported by independent biological libraries, whereas gene-level summaries and partitioned-bin designs require correlation-aware aggregation or joint modelling that the present implementation provides diagnostically rather than generatively. We report this boundary because complex pooled designs make it consequential, not because it is unique to the beta-binomial model.

## 1 Introduction

Pooled CRISPR screens increasingly combine multiple design dimensions, including time courses, ordered FACS fractions, donor cohorts, dose series, and factorial treatment arms, within a single environment. Their scientific quantities of interest are slopes, adjusted associations, and interactions. Reducing these experiments to repeated high-versus-low contrasts discards intermediate observations and prevents one effect from being estimated conditional on the remaining design variables.

The count denominator is central to this problem. RNA-sequencing methods generally model marginal counts with negative-binomial distributions and represent library size through normalization or an offset, which is appropriate when no natural total is available for a gene’s count to be conditioned on [1–5]. In a pooled screen, however, each guide count is observed together with the total number of mapped guide reads in that library, and conditional on this total, guide abundance is a proportion rather than an unconstrained count. The beta-binomial model represents that sampling structure directly, separating depth-dependent sequencing variation from heterogeneity among biological libraries. The original CB^2^ study found better goodness of fit for beta-binomial than negative-binomial models in pooled CRISPR counts [6].

CB^2^ applies this principle to two independent groups, whereas modern screen designs require a general coefficient model. Beta-binomial regression is available in corncob and statistical packages including aod, VGAM, and gamlss [7–10], but these tools do not provide a CRISPR-oriented workflow that preserves immutable full-library totals, fits guide-level design matrices, and connects the result to established screen outputs. Conversely, negative-binomial screen and RNA-sequencing methods, among them MAGeCK-MLE [11], edgeR-QL [5], DESeq2 [3], and limma–voom [4], already accept an arbitrary design matrix, and the comparisons reported below confirm that they do so competitively. The open question is therefore not whether a multivariable screen analysis is possible, but whether the library-total-conditional sampling model—the one component that CB^2^ showed to fit pooled counts better than a marginal negative-binomial model—can be carried into that setting, and what doing so does and does not buy. Specialized methods such as Chronos and Waterbear model population dynamics or joint FACS-bin membership, respectively, but are tied to those experimental structures [12, 13].

We developed BARCS (Beta-binomial Analysis and Regression for CRISPR Screens) to answer that question. BARCS keeps the library-total-conditional beta-binomial sampling model and replaces two group means with a regression design matrix. A single fit can estimate time, donor-adjusted abundance, treatment, or interaction effects. Independent biological libraries, rather than individual reads, determine the residual degrees of freedom. This generalizes the range of questions CB^2^ can address; it does not require the two implementations to share the same estimator or finite-sample test.

We evaluate the extension in increasing order of design complexity and decreasing order of experimental control. A three-time-point Cas13 screen tests longitudinal coefficients across four replicate-complete cell lines. An IL2RA screen tests a donor-adjusted ordered phenotype against independently validated regulators and held-out non-targeting guides. Genome-scale and factorial simulations then isolate dispersion moderation and denominator choice against a known ground truth, including a null-calibration grid that identifies within-gene guide correlation, rather than dispersion alone, as a source of gene-level miscalibration. Finally, an external audit examines how pre-fit subsetting can change the effective library total and an apparent false-discovery result. Together, these analyses assess where the multivariable beta-binomial extension improves on pairwise comparisons, where it matches rather than exceeds existing negative-binomial methods, and where its calibration still requires independent validation.

## 2 Results

BARCS extends the CB^2^ sampling model, which conditions each guide count on the corresponding full-library total to separate sequencing variation from between-library heterogeneity [6]. BARCS retains this sampling principle while replacing the two-group mean comparison with a regression coefficient, allowing time, dose, donor, batch, and interaction effects to be estimated jointly rather than reduced to pairwise contrasts. We evaluate this extension in longitudinal Cas13 and donor-adjusted IL2RA screens, use simulations to isolate the contributions of dispersion moderation and denominator choice, and examine the effect of preprocessing in an external false-discovery audit. Endpoint, serial-harvest, copy-number, and multi-condition follow-up analyses are reported in the Supplement. BARCS therefore broadens the questions that can be addressed with a beta-binomial screen model; it is not numerically equivalent to the legacy CB^2^ statistic, which uses different dispersion, weighting, and degrees-of-freedom calculations.

### 2.1 The intermediate Cas13 time point improves recovery, whereas control scaling is method-agnostic

The Cas13 fitness screens of Liang et al. [14] contain measurements at days 0, 7, and 14. K562 was excluded because one day-0 replicate was not deposited. In HAP1, HEK293FT, MDA-MB-231, and THP1, BARCS, official MAGeCK-MLE [11], edgeR-QL [5], DESeq2 [3], and limma–voom [4] estimated the same continuous time slope with a replicate block and two-sided tests.

The deposited values had already undergone median-of-ratios normalization, ComBat correction, and replicate-outlier processing. We rounded them once to the nearest pseudo-count and supplied the identical matrix to every method. Accordingly, this is a processed-count sensitivity analysis, not a likelihood-faithful comparison of beta-binomial and negative-binomial sampling. The same caution about processing-dependent effective depth raised by the external audit therefore applies here.

The design ablation directly tested whether the middle time point contributed information. Relative to a BARCS fit using only days 0 and 14, the three-time-point fit increased macro-average precision from 0.832 to 0.838 and directional essential-gene recall at FDR 0.10 from 0.538 to 0.600. Recall at an empirical 5% proxy-null false-positive rate increased from 0.858 to 0.867. The proxy-null *p* < 0.05 rate changed from 0.038 to 0.039. Thus day 7 improved essential-gene recovery modestly, while leaving some calibration measures unchanged. Both proxy-null quantities depend on a definition that is itself imperfect. Cell-line-specific nulls are targeted lncRNAs with TPM equal to zero in both available expression assays, so limited expression sensitivity and RfxCas13d collateral activity can each place a genuinely fitness-relevant target in the null set. Because such contamination rewards conservative methods, the proxy-null rates reported here and in the comparison below are diagnostics rather than validated type-I errors.

For the head-to-head calibration analysis, the same 1,000 non-targeting guides were assigned identically to 156–159 pseudo-genes per cell line, with guide counts sampled from the target-gene distribution. The same tail-scaling rule was then applied to each method’s native pseudo-gene summary. Mean cell-line-specific absolute calibration error was 0.0207 for BARCS, 0.0198 for DESeq2, 0.0181 for edgeR-QL, 0.0178 for limma–voom, and 0.0171 for MAGeCK-MLE. Thus aggregation-aware calibration removed the earlier MAGeCK-MLE separation: control scaling is a method-agnostic operation, not a beta-binomial advantage. However, this calibration parity did not carry over to ranking: alternative methods ranked known essential genes more strongly. Average precision was 0.838 for BARCS and 0.874–0.877 for the four alternatives; directional recall at gene FDR 0.10 was 0.600 versus 0.738–0.767. At a matched empirical 5% proxy-null rate, recall was 0.867 versus 0.879–0.888. Thus, once calibration is assessed at the same aggregation level for every method, BARCS matches its competitors in calibration but not in ranking sensitivity.

The gene-level method comparison and three prespecified guide-level disagreement trajectories are shown together in Figure 1, linking the fitted longitudinal coefficient to the observed three-time-point patterns.

**Figure 1:**
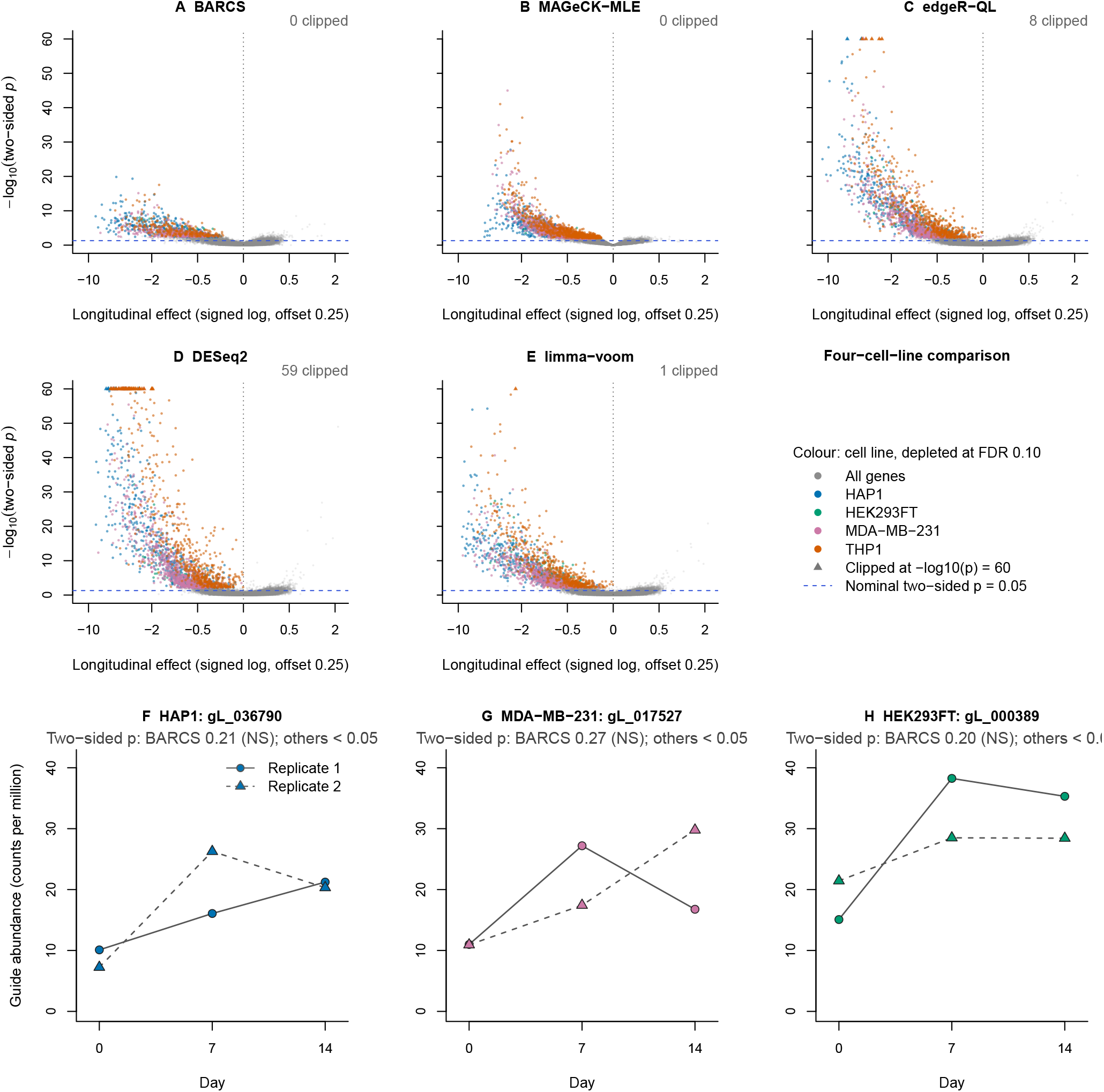
Longitudinal Cas13 sensitivity analysis using days 0, 7, and 14. (A–E) Gene-level results for BARCS, MAGeCK-MLE, edgeR-QL, DESeq2, and limma–voom across the four replicate-complete cell lines. All five panels share one x and y scale and one set of tick marks, so effect sizes are directly comparable between methods. The x axis is the signed logarithmic transform sign (*β*) log_10_ (1 + |*β*|/0.25), labelled at the untransformed effect values. The horizontal line marks two-sided *p* = 0.05; −log_10_(*p*) is clipped at 60, clipped points are triangles, and the count above each panel gives how many points that clip represents. Colour denotes cell line throughout the figure; coloured points are negative-effect genes called at FDR 0.10, and all remaining genes are grey. (F–H) Processed guide abundance in counts per million for three proxy-null disagreements, with markers filled by cell line and replicates distinguished by point shape and line type. Among proxy-null guides, 61 satisfied BARCS *p* ≥ 0.20 with all three guide-level alternatives *p* < 0.05. Within each cell line, the candidate minimizing the largest alternative *p* was identified; the three with the largest ratio of BARCS *p* to that maximum are displayed. Conversely, 1,242 proxy-null guides had BARCS *p* < 0.05 while all three alternatives had *p* ≥ 0.05. The panels therefore illustrate the trajectory structure of one disagreement direction, not its prevalence.

### 2.2 A covariate-adjusted ordered phenotype recovers validated regulators

We next asked whether the design-matrix extension is useful outside a time course. GSE242880 contains an IL2RA protein-abundance screen in primary human T cells, with four ordered FACS bins for each of three donors, 6,000 guides, and 26 regulators validated by individual knockout and flow cytometry [13, 15]. The four bins were assigned prespecified latent normal locations, and BARCS fitted one marker-abundance slope with donor indicators. Official MAGeCK-MLE received the same design. All comparisons below used all four ordered fractions.

This experiment also provides a direct model diagnostic. Bins from one donor partition the same cell pool and are negatively dependent, whereas the guide-level BARCS fit treats their margins independently. Among 593 non-targeting guides, the uncalibrated *p* < 0.05 frequency was 0.133. Because bb_calibrate_controls() estimates its scale from the control tail, evaluating those same guides after fitting would be circular. We therefore used deterministic five-fold cross-fitting: each guide was evaluated with a scale estimated from the other four folds. The held-out *p* < 0.05 rate was 29/593 (0.049). Fold-specific scales ranged from 1.332 to 1.418. A prespecified seeded permutation of the guide-to-fold assignment gave 31/593 (0.052), with scales from 1.332 to 1.417, arguing against structure induced by guide-identifier ordering. The production gene analysis used all controls to estimate one scale; calibration does not change effect estimates or their ranking.

Control-calibrated BARCS recovered 22 of 26 validated regulators with 49 discoveries, compared with 17 of 26 and 72 discoveries for the matched MAGeCK-MLE fit (Table 1). This corresponds to validation panel-hit fractions of 44.9% and 23.6%, respectively; these fractions do not classify the remaining calls as false. Relative to raw four-bin BARCS, control calibration reduced the call set from 127 to 49 while retaining 22 of the 23 validated regulators recovered by the raw analysis. The held-out NTC rate simultaneously moved from 0.133 to 0.049, showing why the calibrated four-bin result is the primary BARCS operating point.

**Table 1:**
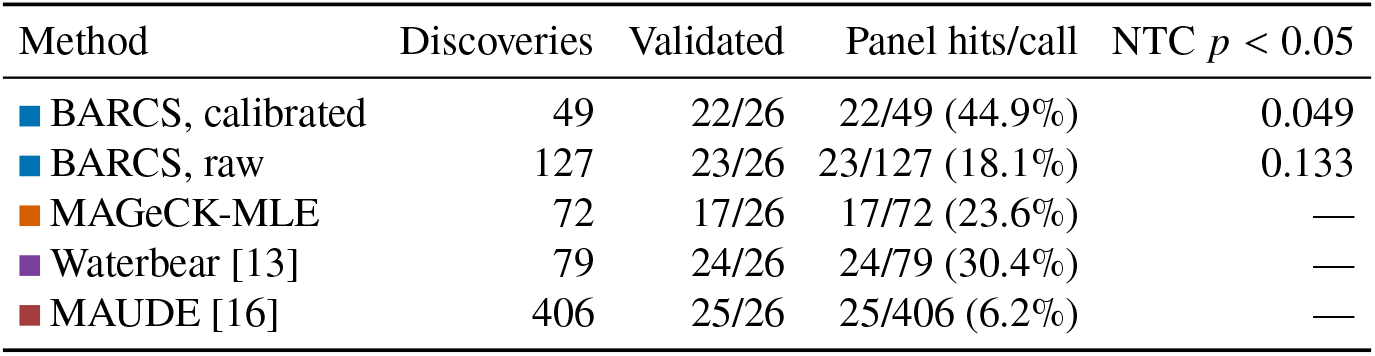
Ordered-bin IL2RA sensitivity analysis. Recovery requires a discovery at each method’s threshold and the experimentally validated direction; all rows use all four FACS bins. The BARCS NTC rate is five-fold held out; it is not measured on the controls used to estimate that fold’s scale. “Validated” is the number of the 26 experimentally validated regulators recovered in the validated direction. “Panel hits/call” divides that number by all method calls. It describes how concentrated a call set is around this limited validation panel; it is not statistical precision or the probability that an individual call is true, because genes outside the panel were not individually validated. An em dash indicates that a comparable guide-level NTC rate was not available.

The specialist models remain important comparators. Waterbear recovered 24 validated regulators and MAUDE recovered 25, but with 79 and 406 discoveries. BARCS therefore offered the strongest recovery-per-discovery ratio among the four-bin models shown, while Waterbear remained the more faithful generative model for correlated sorted bins. Negative controls diagnose the independence violation and calibrate one operating point; they do not make the bin margins independent.

### 2.3 Dispersion moderation improves BARCS in genome-scale simulation

We used CRISPulator to evaluate a FACS design under known truth [17]. At the primary multiplicity of infection (MOI) of 0.20, three prespecified seeds each generated 10,000 genes, five guides per gene, four independent replicates, and low, bulk, and high fractions. All methods received the same counts and design.

The comparison isolates one change within BARCS. BARCS-ST uses the original guide-specific dispersion estimate, whereas BARCS-MOD moderates those estimates before applying the same gene summary. Moderation increased average precision from 0.902 to 0.919 and F1 at gene FDR 0.10 from 0.811 to 0.847, while realized FDP remained nearly unchanged (0.065 and 0.066). This supports dispersion moderation within the common BARCS pipeline.

The moderation gain persisted across requested thresholds at MOI 0.20 and in the secondary MOI-0.30 analysis (Figure 2). Fitting the 50,000 guide-level regressions required 70.1–102.8 seconds per primary seed on the analysis system, providing a direct screen-scale runtime check. MAGeCK-MLE slightly exceeded BARCS-MOD in AUROC and average precision while operating at a much lower realized FDP; CRISPhieRmix operated at a higher FDP (Table 2). Thus the nominal-threshold comparison does not define one universal winner. In the broader 400-gene comparison, edgeR-QL, DESeq2, and limma–voom matched or exceeded BARCS ranking. After matching methods on realized FDP, no method separated from the others above an FDP of 0.005. Ranking and calibration therefore describe different operating characteristics.

**Figure 2:**
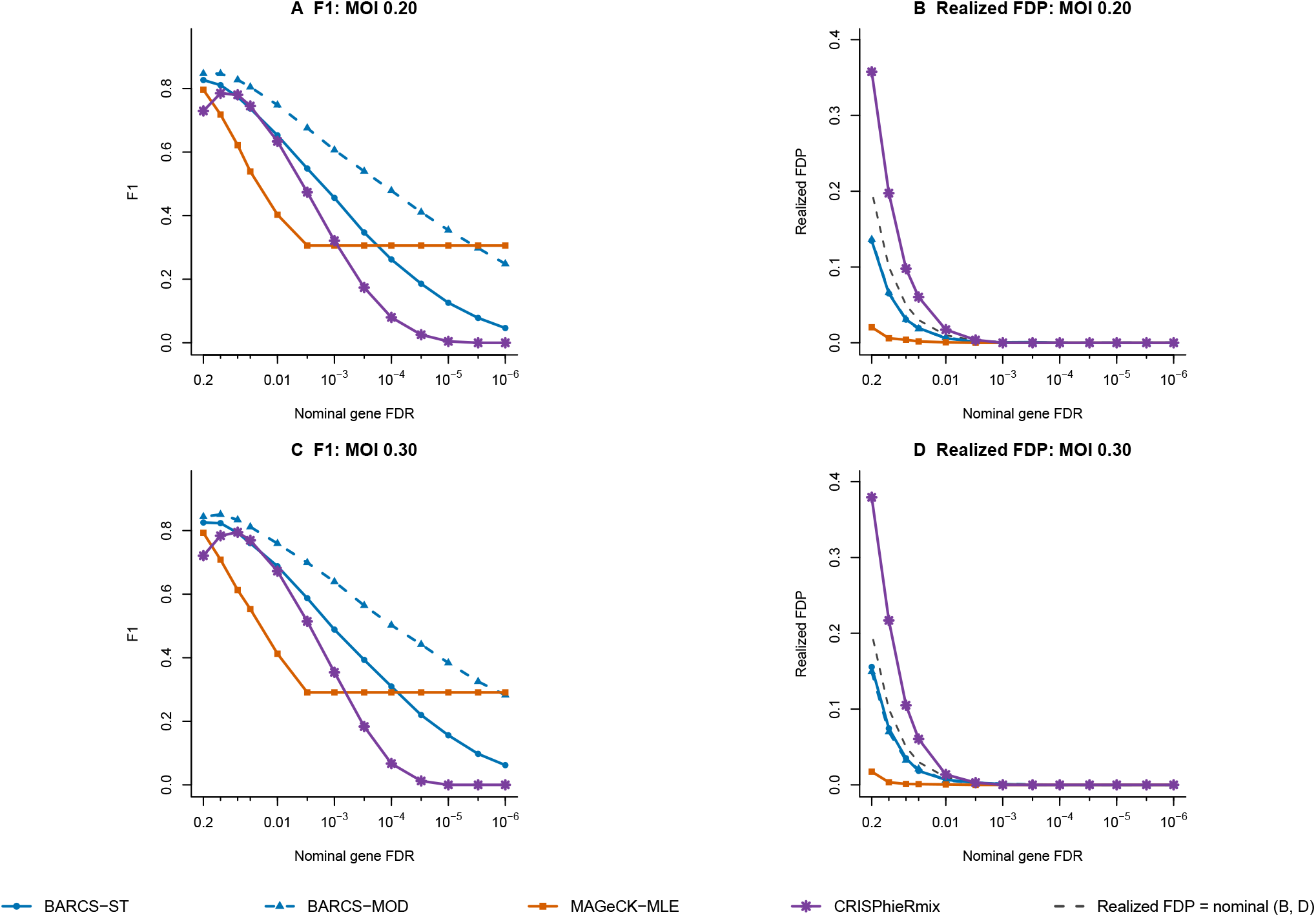
CRISPulator threshold sensitivity. (A,B) MOI 0.20 and (C,D) MOI 0.30. F1 is shown on the left and realized FDP on the right; the dashed diagonal denotes realized FDP equal to the nominal threshold. Both F1 panels share one y scale and both realized-FDP panels share another, and each axis spans the full range of the plotted values, so no curve is cut off at a panel edge. All 13 thresholds carry a minor tick; a non-colliding subset is labelled. BARCS-ST and BARCS-MOD are two settings of one estimator and therefore share a colour, separated by line type and point shape. The shared simulation, gene summary, and thresholds isolate the BARCS-ST versus BARCS-MOD moderation effect.

**Table 2:** Genome-scale CRISPulator comparison for the primary three-bin design at MOI 0.20 (three-seed mean). Each method is evaluated at its nominal gene-level threshold; the resulting operating points are therefore not matched on realized FDP.

| Method | AUROC | Average precision | Realized FDP | F1 |
| --- | --- | --- | --- | --- |
| BARCS-ST | 0.947 | 0.902 | 0.065 | 0.811 |
| BARCS-MOD | 0.954 | 0.919 | 0.066 | 0.847 |
| MAGeCK-MLE | 0.957 | 0.921 | 0.006 | 0.718 |
| CRISPhieRmix | 0.929 | 0.867 | 0.197 | 0.785 |

### 2.4 Control denominators recover interactions under composition shifts

We next used simCRISPR to generate a factorial knockout-by-treatment screen with a known guide-level interaction [18]. Each of three seeds contained 2,000 sgRNAs and three independent replicates per factorial arm. Non-targeting guides had a true interaction of zero, and safe-harbor guides separated the effect of cutting from a targeted gene effect. Each control class was split so that normalization and calibration guides were never used for evaluation.

The full-library denominator preserved effect ranking but shifted the true-zero guides because widespread depletion increased every surviving guide’s library share. Their median fitted interaction was 0.184 (interquartile range 0.136–0.231), whereas the corresponding medians were 0.004 and 0.030 for the non-targeting and safe-harbor denominators. The control-scale factors were 1.99 and 2.24 for unmoderated and moderated full-library fits, compared with 1.00–1.07 for the control-denominator fits. The three-seed means in Table 3 are not representative of a typical run because one library-denominator seed collapsed. BARCS-ST F1 was 0.039, 0.709, and 0.716 across the three seeds; BARCS-MOD F1 was 0.011, 0.680, and 0.754. With the non-targeting denominator, the corresponding values were 0.798, 0.924, and 0.889 for BARCS-ST and 0.899, 0.933, and 0.920 for BARCS-MOD. Thus the control denominator improved every seed, but the magnitude was heterogeneous: in the two noncollapsed runs, the contrast was approximately 0.71 versus 0.91 rather than a near doubling. The held-out null rates remained below the nominal 0.10 in all cases.

**Table 3:** Interaction recovery in simCRISPR (three-seed mean with standard deviation in parentheses). Recall and F1 use targeting guides with |interaction| ≥ 0.2 as positives. Null and safe-harbor rates are held-out call fractions at FDR 0.10.

| Method | Denominator | Dir. recall | F1 | Null rate | Safe-harbor |
| --- | --- | --- | --- | --- | --- |
| BARCS-ST | Library | 0.376 (0.308) | 0.488 (0.389) | 0.004 (0.008) | 0.002 (0.003) |
| BARCS-MOD | Library | 0.375 (0.323) | 0.481 (0.409) | 0.000 (0.000) | 0.000 (0.000) |
| BARCS-ST | Non-targeting | 0.769 (0.094) | 0.870 (0.065) | 0.024 (0.020) | 0.070 (0.025) |
| BARCS-MOD | Non-targeting | 0.838 (0.025) | 0.917 (0.017) | 0.040 (0.040) | 0.080 (0.038) |
| BARCS-ST | Safe-harbor | 0.762 (0.087) | 0.864 (0.061) | 0.042 (0.027) | 0.042 (0.008) |
| BARCS-MOD | Safe-harbor | 0.830 (0.019) | 0.914 (0.017) | 0.027 (0.023) | 0.035 (0.005) |
| MAGeCK-MLE | Non-targeting | 0.003 (0.001) | 0.006 (0.002) | 0.000 (0.000) | 0.000 (0.000) |

Figure 3 shows the composition-induced null shift and the corresponding threshold curves. The two control denominators performed similarly overall but differed on cutting artifacts. Safe-harbor guides were called at 0.080 after non-targeting normalization and 0.035 after safe-harbor normalization. Non-targeting controls are therefore efficient for detecting targeted interactions, whereas safe-harbor controls better absorb the shared cutting response. MAGeCK-MLE is shown only as a reference: the simulated truth is guide-level, leaving no within-gene replication for its native gene model. Accordingly, this benchmark is a denominator and moderation ablation within BARCS, not evidence that the beta-binomial sampling model outperforms a valid interaction comparator. A method-neutral comparison requires a simulation with several independently simulated guides per gene and a common gene-level interaction estimand.

**Figure 3:**
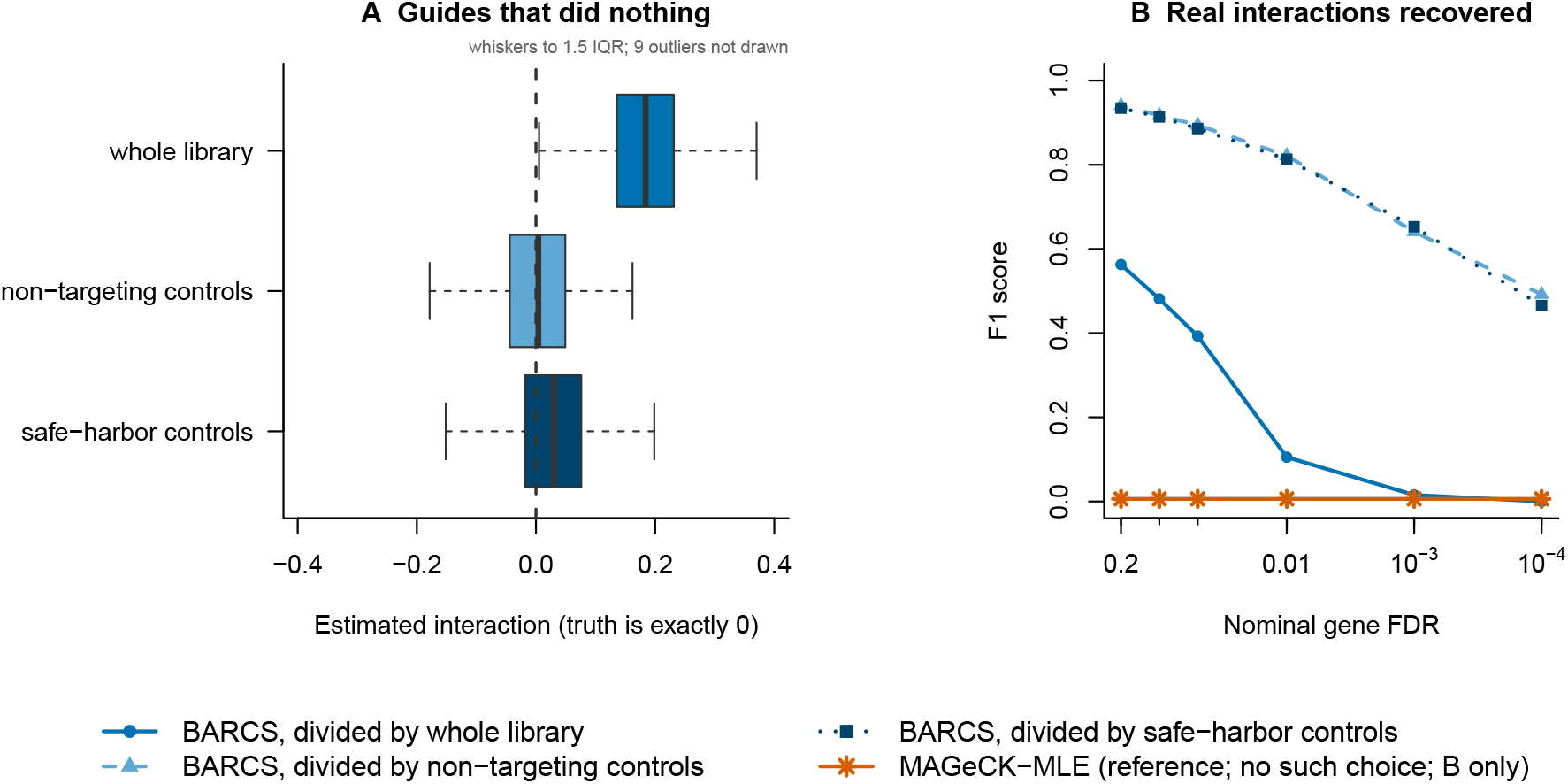
Denominator choice controls interaction recovery. (A) Estimated interactions among true-zero guides. A full-library denominator induces a positive shift; either control denominator removes it. Boxes span the interquartile range, whiskers extend to 1.5 times it, and the number of observations beyond the whiskers is stated in the panel rather than plotted. The value axis spans the full whisker range, so no whisker is cut off at the panel edge. (B) F1 across nominal FDR thresholds, with a minor tick at every threshold and a non-colliding subset labelled. Control denominators retain substantially more interaction power. The three denominators are one estimator run three times and therefore share a hue, separated by lightness, line type, and point shape; MAGeCK-MLE carries its own colour, consistent with the other figures, and appears in (B) only.

### 2.5 Denominator correction removes one artifact but does not resolve calibration

The recent false-discovery manuscript by Dempster et al. [19] raised the concern that CB^2^, and beta-binomial CRISPR analysis more broadly, could be anti-conservative. Because one of the present authors (H.-H.J.) also developed CB^2^, this reanalysis is disclosed as an audit of that author’s prior method rather than an independent adjudication. We revisited its deposited benchmark [20] to distinguish a property of the sampling model from a consequence of the analysis contract. The benchmark compared CB^2^, MAGeCK [11], JACKS [21], DrugZ [22], and Chronos. We separated two analyses that answer different questions. For Avana, we re-evaluated the published CB^2^ analysis [6] after restoring the full-library denominator; BARCS was not fitted because this audit isolates the denominator used by CB^2^. For PSN1 and DeWeirdt, we retained the deposited CB^2^ results and ran BARCS on the deposited normalized tables. In the reported Avana analysis, restriction to null genes before fitting changed the column totals. Reusing the full-library CB^2^ probabilities for the same genes reduced the number of FDR 0.10 discoveries from 152 to zero. This shows that the specific 152-call result depended on the denominator used for fitting; it does not refute the broader claim that beta-binomial analyses can be anti-conservative.

The audit results in Table 4 show that the PSN1 analysis can pool one guide dispersion across the design to avoid the extreme loss of power produced by separate two-replicate variance estimates while retaining a near-nominal guide null. In the DeWeirdt screens [23], however, BARCS recovered only MCL1 under A-1331852; 37 of its 38 calls came from the OVCAR8–olaparib comparison without recovering BRCA1 or BRCA2. A global treatment-associated quality shift cannot be separated from treatment with only two replicates per arm.

**Table 4:** Expected and observed behavior in the external false-discovery audit. Reported results and deposited inputs follow Dempster et al. [19] and the accompanying data release [20]. For an all-null benchmark, a well-calibrated analysis should produce approximately uniform null *p*-values, about 5% below 0.05, and few or no calls after FDR control. For DeWeirdt, “known” denotes 28 curated condition–gene combinations [23], not a complete truth set; therefore no exact number of expected calls can be specified.

| Benchmark | Expected if the analysis is well behaved | Observed result | Interpretation |
| --- | --- | --- | --- |
| Avana null-only | Approximately 5% of null $p$-values below 0.05; few or no calls at FDR 0.10 | Reported CB<sup>2</sup>: 152 calls at FDR 0.10. Full-total CB<sup>2</sup>: 0 calls. BARCS was not run | The 152-call result was denominator-dependent; the residual null rate of 0.145 leaves the broader miscalibration concern unresolved |
| PSN1 null split | Held-out guide $p < 0.05$ rate near 0.05, control scale near 1, and few or no gene calls | Reported CB<sup>2</sup>: 0 gene calls. BARCS: 0 gene calls; guide $p < 0.05$ rates 0.0481 raw and 0.0465 scaled; control scale 1.017 | This result is consistent with the null expectation, but one benchmark does not establish general calibration |
| DeWeirdt differential screens | Known interactions enriched among calls, with no comparison dominating solely because of a quality artifact | Reported CB<sup>2</sup>: 0 calls and 0/28 known recovered. BARCS: 38 calls and 1/28 known recovered; 37/38 calls arose in one comparison | Poor known-interaction recovery and one-comparison concentration suggest comparison-specific confounding |

This reanalysis therefore narrows, but does not refute, the concern raised by Dempster et al. [19]. It shows that pre-fit guide subsetting and common normalization can alter the effective depth seen by a library-total-conditional model, so the reported 152-call example should not be attributed to the sampling model alone. At the same time, zero Avana discoveries after restoring the full total do not demonstrate nominal calibration: the largest cell-line-specific *p* < 0.05 rate remained 0.145, and the DeWeirdt analysis remained unstable. Comparator studies must preserve full-library totals, separate calibration guides from evaluation guides, and use blocked permutations or independent validation for differential screens.

## 3 Methods

### 3.1 Model and inference

For guide *g* in library *i*, BARCS models the count conditional on the unfiltered mapped-guide total:

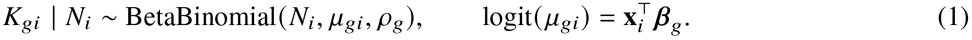

The coefficient model in Equation (1) allows the design vector to contain time, dose, donor, batch, treatment, and interaction terms. Coefficients are estimated by beta-binomial weighted logistic regression. A prespecified coefficient or contrast is divided by its model-based standard error and compared with a Student *t* distribution using residual sample degrees of freedom. All reported tests are two-sided, and guide- or gene-level probabilities are adjusted by Benjamini–Hochberg. The guide-specific overdispersion *ρ*_*g*_ is estimated by solving the Pearson equation at the fitted mean, truncated at zero when the counts are not overdispersed, and the coefficient and dispersion updates are alternated to convergence. BARCS-ST reports these unmoderated statistics. BARCS-MOD additionally shrinks each guide’s fitted variance toward a trend in log mean abundance by an empirical-Bayes step that changes standard errors and residual degrees of freedom but not the coefficient estimates; the real-data benchmarks use unmoderated BARCS. The full estimating equations are given in the Supplement.

### 3.2 Gene-level aggregation and control calibration

Guide coefficients are combined into a gene-level statistic by a prespecified directional Stouffer summary: each guide’s two-sided probability is converted to a normal deviate carrying the sign of its coefficient, the deviates for a gene are combined by the Stouffer rule, and the result is returned to a two-sided gene probability, which is then adjusted by Benjamini–Hochberg. The reported gene effect is the median guide coefficient.

Control calibration rescales a reported statistic without altering the coefficient estimates or their ranking. Given a set of negative-control features, bb_calibrate_controls() maps the 95th percentile of the absolute signed-normal control statistic onto the two-sided standard-normal 0.05 cutoff and divides by the resulting scale, with scales below one truncated at one so that the adjustment can only be conservative. Two properties matter for interpretation. First, the scale must be estimated at the same aggregation level as the statistic it corrects, because a scale learned from individual control guides does not transfer to multi-guide gene statistics; non-targeting guides are therefore grouped into pseudo-genes matching the target-gene guide-count distribution before the rule is applied to a gene-level summary. Second, the controls used to estimate the scale cannot also be used to evaluate it. Wherever a control-based error rate is reported, the scale is obtained by deterministic five-fold cross-fitting and each fold is evaluated with the scale estimated from the other four.

#### Validity of gene-level probabilities

The directional Stouffer summary treats the guides of a gene as independent sources of evidence. Guides targeting the same gene are generally positively correlated—through a shared target effect, a shared cutting response, copy-number-associated toxicity, clonal structure, or unmodeled library composition shifts—and under positive correlation the combined statistic is anti-conservative; the null-calibration grid in the Supplement quantifies the resulting gene-level error. Gene-level probabilities are therefore reported as empirically calibrated ranking statistics rather than as tests carrying a nominal false-discovery guarantee. Guide-level coefficients, their standard errors, and their ordering are unaffected by this limitation, and gene-level inference is recovered when the gene-level null is established by permutation that preserves the within-gene guide structure and the experimental blocking.

### 3.3 Denominator choice defines the estimand

The two denominators available in BARCS answer different questions and are not interchangeable. Under the full-library total, *β* is the change in a guide’s share of the sequenced library. This quantity is always well defined, but it coincides with the change in absolute guide abundance only when the overall library composition is stable across the design; when a large fraction of guides deplete, every surviving guide’s share rises and true-null guides acquire a positive shift (Figure 3). Under a control-set denominator, *β* is the change relative to that control set, and it identifies the absolute effect under the assumption that the control set is invariant across design levels. That assumption should be checked rather than assumed, by comparing the control set’s share of the library across design levels. Non-targeting controls are the efficient choice when the quantity of interest is a targeted effect; safe-harbor controls additionally absorb the shared cutting response and are preferred in Cas9 knockout screens with widespread depletion.

### 3.4 Longitudinal Cas13 analysis

For HAP1, HEK293FT, MDA-MB-231, and THP1, days 0, 7, and 14 were fitted jointly with continuous time and a replicate indicator. K562 was excluded because one day-0 replicate was absent. BARCS, MAGeCK-MLE, edgeR-QL, DESeq2, and limma–voom received the identical rounded matrix and tested the same time slope. Non-targeting guides were grouped into pseudo-genes matching the target-gene guide-count distribution, and the same post-aggregation control scaling rule was applied to all five methods. A BARCS endpoint fit using only days 0 and 14 tested the value of the intermediate time point. Known essential genes were positives; targeted lncRNAs with TPM equal to zero in both available expression assays were proxy nulls. That definition is imperfect in a direction that favours conservative methods, so the proxy-null rates are diagnostics rather than validated type-I errors. Because the deposited values were already normalized and ComBat-corrected, this analysis was treated as a processed-count sensitivity study.

### 3.5 Ordered-bin IL2RA analysis

The IL2RA screen comprised four ordered FACS bins from each of three donors. BARCS fitted one ordered-bin score with donor indicators, and MAGeCK-MLE received the same design. Of 593 non-targeting guides, four folds estimated the control scale and the omitted fold evaluated it; this was repeated across five deterministic folds. A seeded permutation of fold assignments checked that identifier ordering did not determine the result. Method discoveries were compared with 26 regulators validated by individual knockout and flow cytometry. Waterbear and MAUDE results were taken from their published analyses of the same experiment.

### 3.6 Simulation benchmarks

CRISPulator generated three 10,000-gene FACS screens with five guides per gene and four independent replicates. BARCS-ST and BARCS-MOD differed only by guide-dispersion moderation. simCRISPR generated three factorial knockout-by-treatment screens with 2,000 sgRNAs and three replicates per arm. Its known interaction effects allowed direct comparison of full-library, non-targeting, and safe-harbor denominators. Control guides used for normalization and calibration were held out from scoring.

### 3.7 External and supplementary analyses

The external false-discovery audit re-evaluated the deposited Avana CB^2^ probabilities using full-library totals and ran BARCS on the deposited PSN1 and DeWeirdt normalized tables. Supplementary analyses include the serial-harvest HT-29 screen, a four-coefficient GSE70038 screen with method-specific pathway enrichment, and the Sanson A375 endpoint and copy-number audit. The Supplement reports their complete preprocessing, design, and scoring definitions in the same order as the corresponding results.

## 4 Discussion and conclusion

BARCS extends a CRISPR-specific beta-binomial sampling principle from two groups to arbitrary design matrices. The contribution is the coefficient model: time, donor, treatment, and interaction effects can be estimated while each guide remains conditional on its full mapped-guide total. The Liang ablation provides direct, if modest, evidence for the longitudinal extension. Adding day 7 increased average precision from 0.832 to 0.838 and FDR-0.10 essential-gene recall from 0.538 to 0.600 relative to an endpoint fit, while the associated proxy-null rate was essentially unchanged (0.038 versus 0.039).

The like-for-like Liang reanalysis changes the calibration interpretation. All five methods received the same non-targeting-control scaling after the controls were grouped to match the target-gene guide-count distribution. This makes the diagnostic aggregation-aware rather than transferring a one-guide scale to multi-guide gene statistics. Mean absolute calibration errors were similar (0.0171–0.0207), with MAGeCK-MLE numerically closest to nominal and BARCS lower in ranking and FDR recall. Control scaling remains a general calibration device rather than evidence for one count likelihood. Moreover, the matrix was normalized, ComBat-corrected, and rounded before fitting. The result supports a practical coefficient analysis of that processed matrix, not a distributional claim that beta-binomial regression dominates negative-binomial regression.

The IL2RA analysis gives a complementary sensitivity–precision tradeoff. The calibrated four-bin BARCS fit recovered 22 validated regulators with 49 calls, compared with 17 validated regulators among 72 calls for the matched four-bin MAGeCK-MLE fit. Validated regulators therefore comprised 44.9% and 23.6% of the respective call sets, but these are panel-hit fractions rather than estimates of positive predictive value: genes outside the limited validation panel were not individually tested and should not be labeled false. Cross-fitted non-targeting controls reduced the BARCS guide-level *p* < 0.05 rate from 0.133 to 0.049, and a permuted fold assignment gave 0.052. These diagnostics address the operating point; they do not remove the dependence among FACS bins drawn from the same donor.

The simulations identify where BARCS still needs caution. Dispersion moderation improved genome-scale F1, while MAGeCK-MLE slightly exceeded BARCS-MOD in AUROC and average precision at a substantially lower realized FDP. In the interaction simulation, a control denominator improved all three seeds, but one full-library run collapsed; among the other two runs, F1 increased from approximately 0.71 to 0.91. This is a heterogeneous within-BARCS denominator ablation, not a valid between-method comparison. The expanded null grid did not reproduce the continuous-dose failure by increasing dispersion alone under independent guides. Instead, the focused correlated-guide arm isolated the gene combiner: guide-level error remained near or below nominal while Stouffer gene-level error reached 0.108–0.262. Aggregation-matched held-out scaling reduced but did not eliminate that inflation. Guide dependence must therefore be represented in calibration and in future hierarchical gene-level inference.

The external audit corrects one benchmark result but does not refute the broader concern about beta-binomial anti-conservatism. The reported 152 CB^2^ Avana null discoveries did not survive restoration of the full-library denominator, demonstrating that pre-fit subsetting changed the statistical input. This supports the denominator-corrected CB^2^ result and the same immutable-total contract used by BARCS, but it is not evidence of nominal beta-binomial calibration. The largest Avana null *p* < 0.05 rate remained 0.145, and BARCS recovered only one of 28 curated DeWeirdt interactions; both findings leave the broader calibration concern intact.

The defensible conclusion is therefore narrower than a universal method ranking. BARCS supplies a general beta-binomial coefficient model for multivariable CRISPR screens, and the longitudinal and ordered-bin analyses show the sensitivity that additional design columns can recover. Confirmatory use requires independent biological libraries, likelihood-compatible counts, and calibration evaluated on held-out, cross-fitted, or independent controls matched to the aggregation level of the reported statistic. Repeated harvests from one culture and bins partitioned from one donor require blocked, repeated-measures, or joint-compositional models beyond the current independent-library fit. These conditions define where the method is useful and where its probabilities should remain exploratory.

## Data availability

Every dataset analysed in this study is publicly available and no new primary data were generated. The longitudinal Cas13 fitness screens are the deposited supplementary tables of Liang et al. [14]; the corresponding raw sequencing reads are in the NCBI Sequence Read Archive under BioProject accession PRJNA1344834, for which a restartable read-processing pipeline is included with the analysis code. The ordered-bin IL2RA screen is in the NCBI Gene Expression Omnibus under accession GSE242880 [13], and the 26 individually validated regulators used as the recovery panel are those reported by Freimer et al. [15].

The supplementary benchmarks reuse three further public screens. The longitudinal HT-29 time course is Data S2 of Tzelepis et al. [24], obtained through the Europe PMC supplementary bundle for PMC5081405. The Brunello A375 endpoint screen is the count table distributed with CB^2^ [6], originally reported by Sanson et al. [25], with the reference essential and nonessential gene sets of Hart et al. [26]. The four-coefficient multivariable screen is in GEO under accession GSE70038 [27]. Copy-number profiles for the two copy-number audits are from DepMap Public 19Q3 for A375 [28] and DepMap Public 20Q2 for HT-29 (ACH-000552) [29].

The external false-discovery audit uses the deposited benchmark inputs of Dempster [20], available on Figshare at https://doi.org/10.6084/m9.figshare.28850873, together with the published CB^2^ Avana analysis [6]; the differential screens re-examined there are those of DeWeirdt et al. [23].

## Code availability

The BARCS implementation is available at https://github.com/jeonglab-bcm/BARCS.

## Acknowledgements

This work was supported by the Chao Endowment, the Huffington Foundation, and the Jan and Dan Duncan Neurological Research Institute and the Data Science Center at Texas Children’s Hospital.

## A Supplementary information

**Detailed methods.** The following sections provide the estimation and benchmark specifications summarized in the main Methods.

### A.1 Estimation by feasible IRLS

The beta-binomial regression can be written hierarchically as

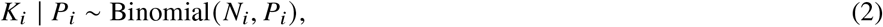

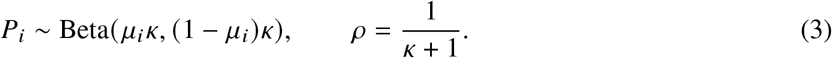

Here *ρ* is the intraclass correlation, or overdispersion, among reads assigned to the same guide and library. For the observed proportion ***Y***_*i*_ = ***K***_*i*_/***N***_*i*_,

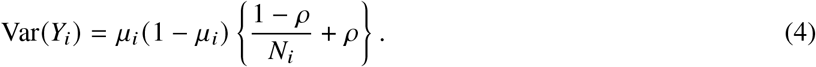

Equation (4) separates sampling variation that decreases with sequencing depth from between-library heterogeneity that does not. This extra-binomial variance formulation and its regression extension follow the line of work in Williams [30], Baggerly et al. [31, 32].

Let *X* be the *m* × *q* full-rank design matrix. Given current values of ***β*** and *ρ*, define

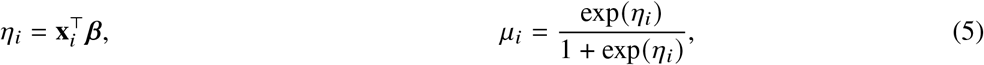

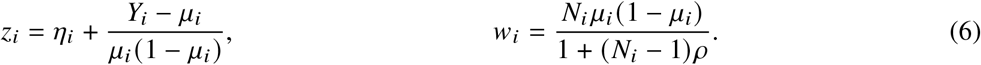

The coefficient update is the weighted least-squares step

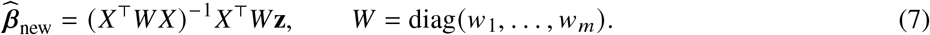

This is weighted least squares on the logistic working response, not on the raw proportions.

For a fixed fitted mean, *ρ* is estimated from the Pearson equation

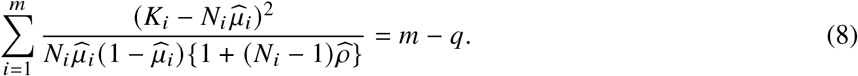

If the left side at *ρ* = 0 is no larger than *m* – *q*, the estimate is truncated at zero rather than claiming negative overdispersion. Equations (6)–(8) are alternated to convergence.

The weight has an important limiting behavior:

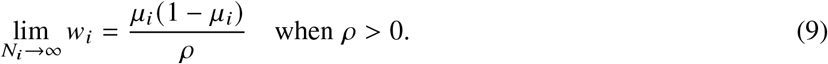

Therefore a deeply sequenced sample cannot acquire unlimited leverage. Once sequencing uncertainty is small, biological heterogeneity sets the information ceiling.

For a continuous design variable *x*_*d*_, exp *β*_*d*_ is the fitted odds ratio for guide abundance per one-unit increase in *x*_*d*_, conditional on the other design columns. Because pooled-screen guide proportions are usually small, *β*_*d*_ also approximates the log abundance ratio per unit. For a contrast, the same interpretation applies to the specified linear combination of coefficients.

Screen-wide weighted cross-products are evaluated with RcppArmadillo [33]; guide fitting remains separable and can be distributed across processor cores.

### A.2 *t*-based inference

At convergence, the model-based covariance estimator is

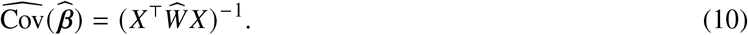

The Pearson equation scales the residual variation to approximately one. If the dispersion estimate reaches its numerical upper boundary, the implementation additionally multiplies Equation (10) by the remaining Pearson scale. Equation (10) is a plug-in, model-based covariance: it conditions on the estimated *ρ* as though it were fixed. It does not propagate uncertainty from estimating a separate dispersion for every guide. Referring the coefficient to *t*_*m*−*q*_ gives a heavier-tailed small-sample reference than a normal approximation, but it is not a formal correction for dispersion-estimation uncertainty. With few independent libraries, calibration must therefore be checked by simulation, phenotype permutation, or prespecified negative controls.

For coefficient *β*_*j*_, the statistic

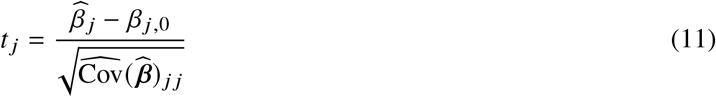

is compared with a Student *t*_*m*−*q*_ distribution. The degrees of freedom are driven by the number of independent sequenced samples, not by the total read count. This is the crucial reason that a library with millions of reads does not create artificial certainty.

More generally, for a prespecified contrast vector **c**,

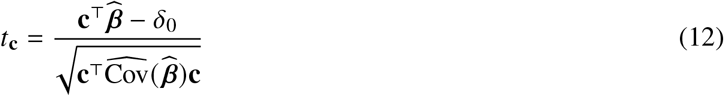

uses the same *m* − *q* degrees of freedom. This covers pairwise group contrasts, average slopes in interaction models, and other one-degree-of-freedom hypotheses. Guide-wise two-sided *p*-values are adjusted using the Benjamini–Hochberg procedure [34].

#### A.2.1 Optional guide-dispersion moderation

BARCS-MOD is an optional empirical-Bayes transformation of the fitted guide statistics, not a separate likelihood. Let *v* = *m* − *q* and let 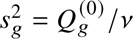 be guide *g*’s Pearson variance inflation under the binomial working model. A lowess trend in log mean CPM gives a guide-specific prior scale 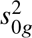, and the scaled-*F* log-moment estimator of Smyth [35] supplies prior degrees of freedom *d*_0_. The moderated inflation is

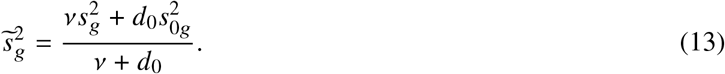

Because the original beta-binomial fit cannot estimate negative overdispersion, its stored standard error corresponds to the fitted inflation 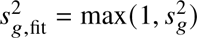. The implementation therefore reports

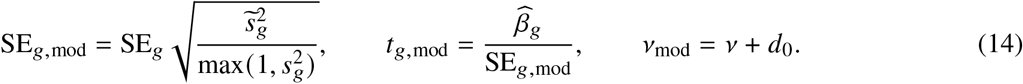

The one-sided pmax truncation in Equation (14) is part of the implementation and is not an estimated scientific constraint. The optional one_way=TRUE variant additionally replaces 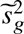 by 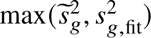, so moderation can only increase a standard error. All reported BARCS-MOD results are simulation ablations; the real-data benchmarks use unmoderated BARCS.

#### A.2.2 From guide statistics to gene statistics

BARCS fits the count model guide by guide. The FACS, simulation, GSE70038, A375, and descriptive HT-29 benchmark scripts then use an explicit directional Stouffer summary, denoted BARCS-ST below, to obtain one gene-level statistic. For the *j*th valid guide targeting gene *g*, the two-sided guide *p*-value is converted back to a signed standard-normal score,

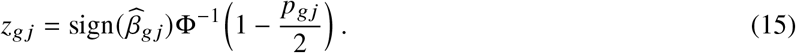

If *m*_*g*_ valid guides target the gene, their combined score and two-sided gene-level *p*-value are

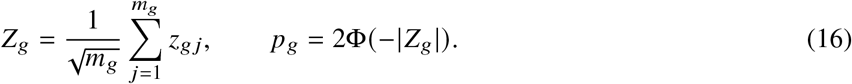

The reported gene effect is the median of the guide coefficient estimates,

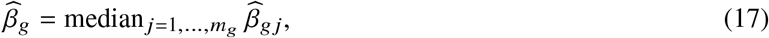

and gene-level false-discovery rates are obtained by applying Benjamini–Hochberg to the collection of *p*_*g*_ values. Consequently, concordant guide directions reinforce one another, discordant directions cancel, and one extreme guide has limited influence on the reported effect.

This aggregation is deliberately transparent, but it is not a native hierarchical gene model. Equations (15)–(16) give equal weight to valid guides and use the independence reference ^√^*m*_*g*_; shared sequence artifacts or other correlation can therefore make the gene statistic anti-conservative. A production extension should estimate guide reliability and within-gene dependence or use a hierarchical effect model. In the biological head-to-head benchmarks, this Stouffer–median procedure is used only for BARCS (and for the deliberately naive binomial comparator). MAGeCK, Chronos, BAGEL2, Waterbear, and MAUDE retain their native or deposited gene-level summaries. The exploratory DeWeirdt false-discovery audit used Fisher aggregation as stated in its Results description and is not pooled with these primary benchmark statistics.

### A.3 Benchmark specifications

#### A.3.1 Liang Cas13 longitudinal fitness benchmark

We analyzed the transcriptome-scale RfxCas13d fitness screens of Liang et al. [14] in HAP1, HEK293FT, MDA-MB-231, and THP1 cells. The library contains 56,322 guides targeting 5,496 lncRNAs, 3,156 protein-coding genes, and 1,000 non-targeting controls. Deposited Table S2 contains guide measurements at days 0, 7, and 14. The previous endpoint analysis selected only days 0 and 14; the present analysis uses all three time points.

The deposited values are fractional after median-of-ratios normalization, ComBat correction, and replicate-outlier processing. They are therefore not literal sequencing counts. BARCS requires integer observations, so each value was rounded once to the nearest pseudo-count and the identical rounded matrix was supplied to every newly fitted method. The mean absolute rounding change was approximately 0.25 and the maximum was less than 0.5 in every cell line. Consequently, this benchmark is a same-input sensitivity analysis rather than a likelihood-faithful raw-count comparison.

BARCS, MAGeCK-MLE, edgeR-QL, DESeq2, and limma–voom fitted a linear longitudinal effect with time scaled from 0 to 1 over 14 days and a replicate indicator. K562 was excluded because one deposited day-0 replicate is absent, rather than fitting an unmatched trajectory without the replicate block. For a like-for-like calibration comparison, non-targeting guides were assigned deterministically to pseudo-genes whose guide counts were sampled from the observed target-gene guide-count distribution in each cell line. The same pseudo-gene assignments were supplied to all five methods. After each method’s native gene summary, the 95th percentile of the absolute signed-normal pseudo-gene statistic was mapped to the two-sided standard-normal 0.05 cutoff, with scales below one truncated at one. This estimates the scale after guide aggregation rather than transferring a one-guide scale to multi-guide genes. Calibration did not alter effect estimates or ranks. All methods used two-sided guide or Wald probabilities. For the four guide-level regression methods, coefficient direction was retained in the signed Stouffer score; the resulting gene probability was two-sided and was adjusted by the Benjamini–Hochberg procedure. Endpoint robust-rank-aggregation summaries [36] were excluded because they do not estimate the three-time-point longitudinal coefficient.

Known-essential protein-coding genes were positives. Cell-line-specific nulls were targeted lncRNAs with TPM equal to zero in both available expression assays. This proxy-null definition is itself imperfect: TPM equal to zero can reflect limited expression sensitivity, and RfxCas13d collateral activity can produce genuine fitness effects from a nominally unexpressed target. Such contamination rewards conservative methods, so the reported null rates are diagnostics rather than validated type-I errors. We reported average precision, essential-control recall at an empirical 5% null false-positive rate, the absolute deviation of the null *p* < 0.05 fraction from 0.05, and directional essential-control recall at gene FDR 0.10. A restartable raw-read pipeline is also provided for BioProject PRJNA1344834 using the stated Cutadapt anchors [37] and Bowtie -v 1 -m 3 –best -q [38].

#### A.3.2 Ordered-bin IL2RA benchmark

The GSE242880 count matrix contains four ordered FACS bins for each of three donors, 6,000 guides, and 593 non-targeting guides [13, 15]. Bin locations were set to the expected values of four equal-probability intervals under a standard normal distribution. BARCS fitted this ordered score with donor indicators. MAGeCK-MLE received the same four-bin design matrix. All reported IL2RA method comparisons used all four ordered fractions.

BARCS guide results were aggregated with the prespecified directional Stouffer summary. For held-out calibration, non-targeting guides were ordered by guide identifier and assigned cyclically to five folds. Each fold was evaluated with the tail scale estimated from the other four folds. The production gene analysis used all controls, whereas the reported non-targeting *p* < 0.05 rate uses only held-out guide predictions. As a structure check, the same cross-fitting calculation was repeated after a seeded permutation of guide identifiers; this changes only fold membership, not the fitted guide coefficients.

Directional recovery was evaluated against 26 regulators validated by individual knockout and flow cytometry. A recovered regulator had to pass the method’s reported threshold and agree with the validated direction. The panel-hit fraction was calculated as recovered validated regulators divided by all discoveries. Because validation was limited to this panel, that fraction is a descriptive measure of call-set concentration, not an estimate of positive predictive value; discoveries outside the panel were not counted as validated but were not assumed to be false. Published Waterbear and MAUDE totals were used only for this recovery comparison; no unavailable non-targeting rate was imputed.

#### A.3.3 CRISPulator FACS benchmark

We used CRISPulator 0.5.1’s documented interfaces for library construction, transfection, FACS selection, and sequencing [17]. The committed Julia environment pins the tagged source revision. For each simulation seed, one CRISPRn guide library was constructed and reused across four independently seeded transfection, sorting, and sequencing replicates. The default CRISPulator phenotype mixture was retained: 75% inactive, 5% negative-control, 10% phenotype-increasing, and 10% phenotype-decreasing genes. The headline screens contained 10,000 genes and five guides per gene, so 50,000 guides, which is the order of a genome-wide library rather than a pilot. The default guide-quality mixture—90% complete-knockout guides and 10% low-activity guides—was retained. Transfection representation was 500 cells per guide, Gaussian phenotype noise was 2, sorting representation was 50 cells per guide, and sequencing depth was 50 reads per guide per sample. Multiplicity of infection was 0.20 for the main analysis and 0.30 for a secondary sensitivity analysis; 0.20 is the point at which the single-integration approximation these guide-level models share is most defensible, and 0.30 stresses it. The three genome-scale benchmark seeds were 20250724 through 20250726. The primary table reports only MOI 0.20; MOI 0.30 appears in the threshold sensitivity figure. The supporting 400-gene analyses use seeds 20250724 through 20250728.

Two BARCS gene statistics are carried through the genome-scale ablation. BARCS-ST is the historical calibrated signed-*z* statistic. BARCS-MOD reuses the identical guide fits after guide-dispersion moderation, so the two differ only in how the guide variance is estimated, not in the gene combiner. Official MAGeCK-MLE 0.5.9.5 and CRISPhieRmix are included in that run, but this restricted set is not treated as an exhaustive comparator panel. MAGeCK-MLE’s ordered marker is affinely mapped onto zero–one so that the low samples of the reference replicate form the all-zero reference row the MAGeCK initializer requires. Because MAGeCK-MLE omits negative-control genes from its gene summary once their sgRNAs are declared, all methods in the genome-scale benchmark are scored on the genes every method returned a finite result for, 9,475–9,517 of 10,000 per run. The negative-control diagnostic is instead taken from each method’s own output, and is therefore unavailable for MAGeCK-MLE.

CRISPulator’s arbitrary percentile ranges were set to 0, 0.25 for low, 0.75, 1 for high, and 0, 1 for bulk. Thus bulk deliberately overlaps the tails and is not a third component of a multinomial partition. An input sample was generated as an independent sequencing draw from the same guide frequency vector as bulk. Low and high received phenotype scores equal to the means of their corresponding standard-normal intervals,

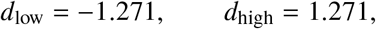

while bulk received *d*_bulk_ = 0. BARCS and MAGeCK-MLE fitted this score with replicate indicators [39]. Tail-only fits removed bulk without changing the low and high counts. BARCS guide tests were scaled with the simulated negative-control guides before directional Stouffer aggregation; MAGeCK-MLE used its native gene-level Wald result. The separate 400-gene, five-seed baseline includes edgeR-QL, DESeq2, and limma–voom in addition to BARCS and MAGeCK-MLE. It is used to prevent the restricted genome-scale set from supporting a general method-ranking claim. CRISPhieRmix [40] was run on guide-level log_2_ fold changes from a DESeq2 Wald fit of the same design [3], which is the input its documentation specifies; DESeq2 therefore appears as a preprocessing step in the 10,000-gene run, while its native result is scored in the separate 400-gene comparison. CRISPhieRmix was run with its bimodal alternative, because this screen contains both phenotype-increasing and phenotype-decreasing genes and the one-sided default would score only one direction. Its gene effect is the mean guide log_2_ fold change and its local false-discovery rate stands in for a *p*-value wherever a ranking is required, since the model reports neither a *p*-value nor a signed gene effect. Reported runtimes cover the primary low–bulk–high model fit and exclude simulation, file input/output, and guide-to-gene aggregation.

Genes from the increasing and decreasing classes were positives. Ranking used the two-sided gene significance score. AUROC and average precision measure active-gene discrimination. Effect recovery is Spearman correlation between the inferred gene effect and CRISPulator’s median theoretical guide phenotype among active genes. Directional recall is the fraction of active genes with gene FDR below 0.10 and an effect sign matching the simulated class. Realized FDP is the fraction of called genes that were inactive, and F1 uses active versus inactive status at the same FDR threshold.

#### A.3.4 simCRISPR interaction benchmark

CRISPulator assigns each guide a single effect and sorts cells on a phenotype, so it cannot generate the design BARCS is aimed at. simCRISPR [18] simulates a factorial screen instead: knockout induction crossed with treatment, propagated over several days of cell growth, with PCR amplification and sequencing applied afterwards. Every sgRNA therefore carries a knockout effect, a treatment effect, and an interaction between the two, and that interaction is a coefficient in a design matrix rather than a contrast of two arms.

Each of three seeds generated 2,000 sgRNAs, of which 300 were non-targeting and 200 were safe-harbor controls, in three independent replicates of each of the four arms, so twelve libraries and eight residual degrees of freedom. Growth was logistic. The alternative exponential model is unbounded: over five days a single guide reached a fifth of the library and library sizes spanned a 175-fold range, which does not represent a pooled screen. Initial library columns were dropped, since they are not part of the factorial contrast. Amplification used the package defaults and each library was sequenced to 500 reads per guide. BARCS fitted logit (*µ*)_*gi*_ = *β*_0_ *β*_1_*k*_*i*_ *β*_2_*t*_*i*_ *β*_3_*k*_*i*_*t*_*i*_ for knockout indicator *k*_*i*_ and treatment indicator *t*_*i*_, and tested *β*_3_.

Three denominators were compared. The library denominator is the full column total, BARCS’s default. The two control denominators hold a chosen control class at a constant share, rescaled to the mean library size and rounded, so that they remain interpretable as library sizes. The two control classes are not interchangeable: safe-harbor guides are cut and carry the DNA-damage response, non-targeting guides are neither. Each control class was split in half, one half normalizing and calibrating and the other held out, so that no guide used to fit the null was also scored against it. MAGeCK-MLE received the same design matrix with an explicit interaction column, the same calibration-half controls, and one sgRNA per gene, because the simulated truth is guide-level.

CRISPhieRmix was not run on these data. It borrows strength across the guides within a gene, whereas simCRISPR assigns every sgRNA an independent interaction, so grouping guides into synthetic genes would impose the structure that the model exploits.

Every targeting guide receives a nonzero interaction, most far too small to resolve, so a magnitude was fixed before scoring: guides with INT 0.2 are the positives and the held-out non-targeting guides, whose interaction is exactly zero, are the negatives. Targeting guides below that magnitude are scored for effect recovery but excluded from the detection metrics. Effect recovery is the Spearman correlation between the estimated interaction coefficient and the simulated one across all targeting guides, which needs no threshold. The held-out control call rate and the safe-harbor call rate are reported separately: the first measures false positives against a ground-truth zero, the second measures how often the cutting response alone is mistaken for an interaction.

#### A.3.5 Longitudinal HT-29 benchmark

Guide counts for the HT-29 time course of Tzelepis et al. [24] were obtained from Data S2 of that article through the Europe PMC supplementary bundle for PMC5081405. The three sequencing columns reported at each of days 7, 10, 13, 16, 19, 22, and 25 were summed to one column per harvest and combined with the pDNA measurement, and guides with fewer than 30 pDNA reads were removed, leaving 86,882 guides targeting 17,995 genes. Library totals were computed before that filter and never recomputed, so the beta-binomial denominator remains the full mapped-read total for each sample; the filter removes 4.2% of guides but only 0.10% of reads, uniformly across samples. BARCS and official MAGeCK-MLE 0.5.9.5 were both fitted to this eight-column matrix under an identical design, in which pDNA is day 0 and the tested predictor is day divided by 25, so the two continuous-time fits differ only in the likelihood. A minimum total count of 30 was applied within the BARCS fit. Because all late samples are serial harvests of one culture, these fits are used as descriptive trajectory scores; their residual-*t* or Wald probabilities are not interpreted as independent-sample inference.

Comparator effects were not recomputed. The joint Chronos, per-endpoint MAGeCK, and per-endpoint BAGEL2 effect matrices were taken as deposited in Chronos Figshare article 14067047 [41], as were the reference essential and nonessential gene lists; the corresponding individual file identifiers are 26548532, 26548541, 30847513, 26548550, and 26548553. The latter two matrices carry one column per day; the day-25 column is used directly, so the comparator is a single observed endpoint at the deepest time point rather than a summary across days.

Two external annotations define the evaluation sets. Positives are the deposited reference essential genes. Negatives are HT-29 genes with DepMap Public 20Q2 expression below 0.5, taken from the ACH-000552 column of Figshare file 26548487, which is the expression source used for the same threshold in the deposited Chronos analysis. The copy-number audit uses the ACH-000552 column of Figshare file 26548484 [29], and correction was performed with MAGeCK 0.5.9.5’s official piecewise –cnv-norm normalizer applied after model fitting. Because that profile was measured independently of the screen, cell-line drift is a limitation of the copy-number analysis.

The deposited workflow records the source identifier and checksum of every downloaded input, executes the official MAGeCK model and copy-number normalizer, and reproduces the tables reported here.

#### A.3.6 GSE70038 multivariable screen

The 64,747-guide count matrix for GSE70038 was downloaded from GEO [27]. Following the Table 5 and Box 3 construction in the MAGeCKFlute protocol [42], the 16 libraries received an intercept for their shared initial condition and four indicators for the GSC0131, GSC0827, NSCCB660, and NSCU5 terminal samples. BARCS used the unfiltered column sum as ***N***_*i*_ and fitted all four coefficients jointly. Official MAGeCK-MLE 0.5.9.5 received the same count matrix and design and used its median normalization [11].

BARCS guide probabilities were combined within each gene by a two-sided directional Stouffer statistic; the gene effect was the median guide coefficient. The complementary analysis selected the 200 most negative finite-effect genes from each method for each coefficient, partitioned the equal-sized lists into shared and method-specific sets, and queried GO Biological Process 2023 and Reactome 2022 through Enrichr [43]. As an external follow-up, one-sided Fisher tests measured enrichment of each exclusive set for the Hart reference-essential genes [26] against the genes testable by both methods.

#### A.3.7 Sanson A375 endpoint and copy-number audit

The Brunello A375 screen contained plasmid and terminal biological libraries with guide-to-gene mappings and the reference-essential and nonessential sets distributed with CB^2^ [25, 26]. BARCS fitted a plasmid-versus-terminal coefficient to each guide using the full mapped-guide totals. Guide evidence was aggregated by a two-sided directional Stouffer statistic, with the median guide coefficient as the gene effect. Official MAGeCK-MLE 0.5.9.5 received the same counts and two-column design and used median normalization [11].

AUROC, average precision, directional recall, and realized false-discovery proportion were evaluated on the common copy-number-complete reference set. Uncertainty in paired AUROC and average-precision differences was estimated by 2,000 stratified bootstrap resamples of genes. Copy-number sensitivity used the A375 profile from DepMap Public 19Q3 [28]. The official MAGeCK piecewise post-fit correction was applied to both methods’ gene-effect vectors; probabilities were not recomputed because the correction changes the effect ranking rather than refitting either likelihood [44].

**Additional analyses.** These analyses extend the main evaluation without changing its primary longitudinal and multivariable claims.

### A.4 Nominal-versus-realized null calibration

We first report the previously specified continuous-dose simulation because it exposes a finite-sample failure that a methods paper should not hide. At a nominal 0.05 threshold, unmoderated BARCS had gene-level type-I error 0.081 and empirical FDP 0.130; moderation reduced FDP to 0.091 but did not repair the marginal type-I error (Table S1). Official MAGeCK-MLE was slightly conservative in this realization. These results show that neither the plug-in beta-binomial statistic nor its moderated form is uniformly calibrated without an external control diagnostic.

**Supplementary Table S1:** Nominal-versus-realized error in the prespecified continuous-dose simulation. Type-I error is the fraction of 160 null genes with *p* < 0.05; empirical FDP is measured among calls at Benjamini–Hochberg FDR 0.05. This single deterministic simulation is a diagnostic, not a uniform operating characteristic.

| Method | Null type I | Power | Calls | Empirical FDP |
| --- | --- | --- | --- | --- |
| Naive binomial $z$ | 0.844 | 1.000 | 174 | 0.770 |
| BARCS unmoderated $t$ | 0.081 | 1.000 | 46 | 0.130 |
| BARCS moderated $t$ | 0.088 | 1.000 | 44 | 0.091 |
| Official MAGeCK-MLE Wald | 0.038 | 0.975 | 41 | 0.049 |

We then simulated all-null beta-binomial screens with *m* ∈ {4, 6, 8, 12} balanced independent libraries, 100 genes, three or five guides per gene, and a library total of 100,000. Mean guide counts were 10, 100, or 1,000. Dispersion was parameterized by (*N* − 1)*ρ* ∈ {0, 5, 60, 180}, corresponding to *ρ* ∈ {0, 0.00005, 0.00060, 0.00180} and spanning the 60–180 range in the continuous-dose simulation. Cells combining mean count 10 with (*N* − 1)*ρ* ≥ 60 were omitted because fewer than 50 guide fits were estimable. Each remaining combination used five prespecified seeds.

The tested group coefficient was zero for every guide. Guide probabilities were combined by the directional Stouffer rule used in the benchmarks. To test calibration after aggregation, odd-numbered null genes estimated a signed-normal 95th-tail scale and even-numbered null genes evaluated it. A focused arm at *m* = 8, mean count 100, and (*N* − 1)*ρ* ∈ {60, 180} generated beta probabilities with a Gaussian copula having within-gene guide correlation 0.4. Table S2 reports ranges across the applicable dispersion, abundance, and sample-size cells. Complete parameter-by-parameter and seed-level values accompany the manuscript.

**Supplementary Table S2:**
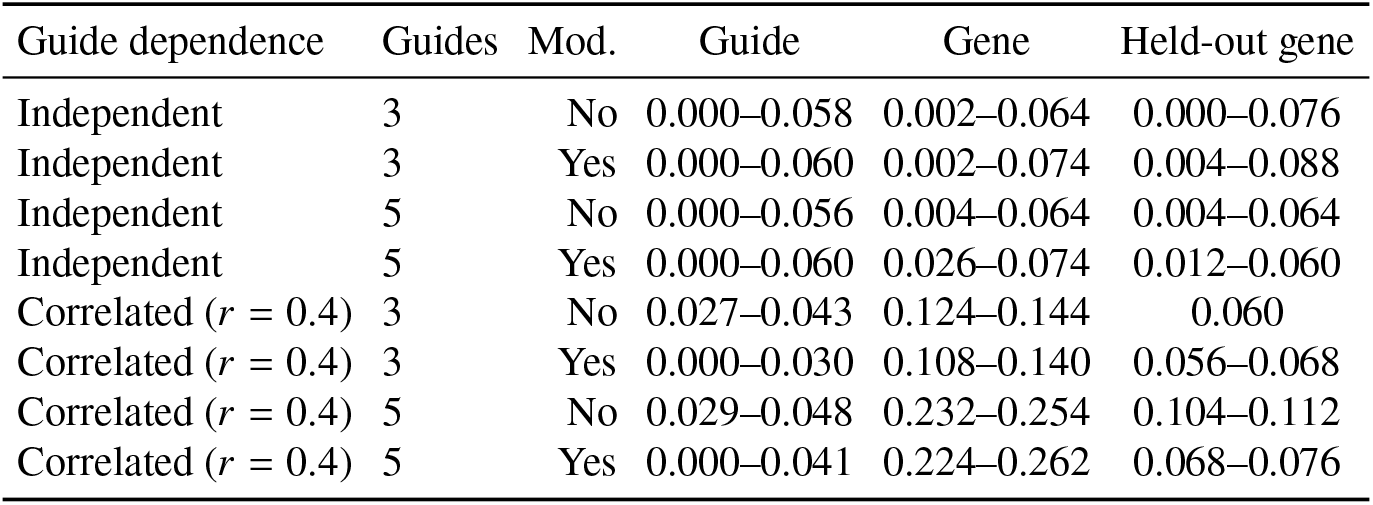
Null calibration before and after gene aggregation. Entries are ranges of five-seed mean rejection rates across the applicable grid cells. The held-out column applies a scale learned from a disjoint half of the null genes after the same aggregation. The target rate is 0.05.

Under independent guides, expanding dispersion through the continuous-dose range did not reproduce the 0.081 gene-level error: the largest five-seed mean was 0.074, and error did not increase monotonically with dispersion. The continuous-dose failure therefore cannot be attributed to dispersion alone. In contrast, within-gene guide dependence left guide-level error near or below 0.05 but raised gene-level error to 0.108–0.262. Aggregation-matched split-control scaling reduced this range to 0.056–0.112, but did not fully repair the five-guide correlated cells. The gene combiner and within-gene dependence are therefore material parts of the calibration problem, and a control scale must be estimated at the same aggregation level as the reported statistic.

Finally, the observed fraction of converged guide fits with *ρ* = 0 was 11/86,840 (0.013%) in HT-29, 0/5,999 in IL2RA, and 1,738/280,541 (0.620%) across the four Liang cell lines. Boundary truncation is therefore common in the deliberately near-binomial grid but rare in the three real-data fits. That observation rules out the boundary as a general explanation for their calibration behavior; limited independent-library degrees of freedom and the plug-in treatment of dispersion remain relevant.

### A.5 HT-29 serial harvests are descriptive, not replicated longitudinal inference

The HT-29 dataset of Tzelepis et al. [24] contains one plasmid pool and guide counts from days 7, 10, 13, 16, 19, 22, and 25 of one passaged culture. The three sequencing columns at each late day were summed because they assay the same harvested population. The seven harvests are likewise repeated measurements of one biological lineage, not seven independent biological units. Referring a time coefficient to *t*_6_ would therefore be pseudoreplication and would violate the independent-library assumption.

We retain the fitted BARCS and official MAGeCK-MLE time coefficients only as descriptive trajectory scores. They received the same eight-column matrix and numeric design. We also report the deposited all-time-point Chronos score and day-25 MAGeCK and BAGEL2 scores [12, 41, 45]. Core essential genes are positives and HT-29 genes with DepMap 20Q2 expression below 0.5 are negatives, matching the deposited evaluation.

**Supplementary Table S3:** Descriptive HT-29 control-gene ranking. The BARCS and MAGeCK time rows treat serial harvests as a numeric trajectory for ranking only; their sample-level *p*-values are not interpreted. Recall_90_ is maximum recall at at least 90% precision, and FP_10%_ counts unexpressed genes in the lowest 10% of all evaluated effects.

| Method | AUROC | PR AUC | NNMD | Recall<sub>90</sub> | FP<sub>10%</sub> |
| --- | --- | --- | --- | --- | --- |
| ■ BARCS time | 0.9786 | 0.9494 | −16.34 | 0.9037 | 7 |
| ■ Official MAGeCK time | 0.9789 | 0.9534 | −17.75 | 0.8972 | 5 |
| ■ Chronos joint | 0.9747 | 0.9356 | −27.88 | 0.8991 | 10 |
| ■ Deposited MAGeCK day 25 | 0.9792 | 0.9428 | −14.83 | 0.9120 | 10 |
| Deposited BAGEL2 day 25 | 0.9698 | 0.9388 | −12.06 | 0.8630 | 4 |

In Table S3, the day-25 MAGeCK row has the highest AUROC and Recall_90_ in the deposited comparison. The full trajectory therefore provides no measured advantage over the endpoint on these two ranking metrics. Official continuous-time MAGeCK has the highest PR AUC, Chronos has the most negative NNMD, and BAGEL2 has the fewest unexpressed genes in the 10% tail. These differences illustrate that the metrics reward different score properties; none supplies the missing biological replication.

How a deposited time course is summarized also matters. Taking the median MAGeCK effect across all seven days instead of the day-25 column lowers AUROC from 0.9792 to 0.9742 and Recall_90_ from 0.9120 to 0.8602. Early harvests carry little depletion, so an across-day median attenuates the endpoint signal. This data-reduction effect is larger than the differences among the trajectory scores and further argues against treating this dataset as inferential evidence for one count model.

### A.6 Copy-number sensitivity in the longitudinal HT-29 screen

We extracted the HT-29 profile (ACH-000552) from DepMap Public 20Q2 [29] and repeated the same copy-number audit used for A375. Before correction, effect–copy-number Spearman correlation among unexpressed genes is already small: 0.072 for BARCS, 0.075 for official continuous-time MAGeCK, 0.048 for deposited Chronos joint effects, 0.057 for the published day-25 MAGeCK effects, and 0.029 for the published day-25 BAGEL2 effects. Applying MAGeCK 0.5.9.5’s official piecewise normalizer makes the association larger in absolute value rather than smaller: 0.145 for BARCS and 0.149 for MAGeCK. Thus the improvement observed in A375 does not generalize automatically.

The piecewise correction estimates one global effect–CNV trend across genes in one cell line and applies it after model fitting. When the nuisance association is weak and biological dependencies are not exchangeable across copy-number regions, the fitted trend can over-correct null genes. Chronos joint is shown without CNV correction because the published procedure requires multiple cell lines to identify common essential genes and was not applied to this single-line experiment [12]. The DepMap profile was measured separately from the original screen, so cell-line drift is an additional limitation. These results support diagnosing copy-number correction rather than applying it by default.

### A.7 Complementary hits in a four-condition screen

GSE70038 contains 64,747 guides in 16 libraries from two glioblastoma stem-cell lines and two neural stem-cell lines [27]. BARCS and MAGeCK-MLE received the same five-column design: one shared initial condition and four cell-line-specific terminal coefficients. This analysis tests whether the multivariable interface estimates all four effects jointly; MAGeCK agreement is not treated as biological ground truth.

To examine the disagreements directly, we selected the 200 most depleted genes from each method for each terminal coefficient. This equal-size rule avoids comparing differently calibrated FDR cutoffs. The lists shared 139–144 genes, leaving 56–61 BARCS-only and the same number of MAGeCK-only genes per cell line (Supplementary Table S4).

**Supplementary Table S4:** Complementary GSE70038 top-200 depletion sets and their strongest Reactome 2022 enrichment. Values in parentheses are Enrichr-adjusted *p*-values.

| Coefficient | Shared | BARCS only | MAGeCK only | Top BARCS-only pathway | Top MAGeCK-only pathway |
| --- | --- | --- | --- | --- | --- |
| GSC0131 | 142 | 58 | 58 | Cell cycle ($1.35 \times 10^{-6}$) | rRNA processing ($1.65 \times 10^{-5}$) |
| GSC0827 | 139 | 61 | 61 | RNA metabolism ($2.77 \times 10^{-9}$) | RNA metabolism ($2.35 \times 10^{-4}$) |
| NSCCB660 | 140 | 60 | 60 | RNA metabolism ($2.10 \times 10^{-4}$) | RNA metabolism ($2.21 \times 10^{-6}$) |
| NSCU5 | 144 | 56 | 56 | RNA metabolism ($3.36 \times 10^{-12}$) | RNA metabolism ($2.80 \times 10^{-13}$) |

Enrichr analysis of the complementary sets used GO Biological Process 2023 and Reactome 2022 [43]. Both methods’ exclusive hits formed coherent essential-process groups. BARCS-only GSC0131 hits emphasized cell cycle regulators, whereas the MAGeCK-only set emphasized rRNA processing; RNA metabolism dominated both methods in the other coefficients. These results provide a biological follow-up to the method disagreement, but not an accuracy ranking: Enrichr uses its library background rather than the genes testable in this screen, and essential-process enrichment is expected in a depletion experiment. Against the independent Hart reference set, 62.3–75.9% of the BARCS-exclusive genes and 59.0–86.2% of the MAGeCK-exclusive genes were essential, with strong enrichment for both methods in every coefficient (one-sided Fisher *p* < 10^−22^). Neither method was consistently higher. The complete overlap and enrichment records are provided with the analysis output.

### A.8 Endpoint benchmark and copy-number audit

We used the Brunello A375 negative-selection screen of Sanson et al. [25], bundled with CB^2^, as a descriptive endpoint benchmark. Reference essential genes are positives and reference nonessential genes are negatives [26]. The common copy-number-complete set contains 1,506 essential and 869 nonessential genes, so its positive prevalence is 63.4%. F1 and recall in this enriched set must therefore be interpreted together with the realized false-discovery proportion.

Copy number is a gene-level property of A375 and is constant across the plasmid and terminal libraries, so it cannot be identified as a sample-level coefficient. We instead applied MAGeCK 0.5.9.5’s official post-fit piecewise correction to both methods’ gene effects using the A375 profile from DepMap Public 19Q3 [28, 44]. This changes effect rankings but does not refit either count likelihood.

**Supplementary Table S5:** Sanson A375 endpoint sensitivity analysis. Recall and gold-set FDP use nominal FDR 0.05 and a negative effect. The last column is Spearman correlation between effect and copy number among reference nonessential genes. No maximum is highlighted because the ranking confidence intervals overlap and the gold-standard set is enriched for positives.

| Method | AUROC | Avg. precision | Recall | Gold-set FDP | CNV corr. |
| --- | --- | --- | --- | --- | --- |
| ■ BARCS | 0.9598 | 0.9786 | 0.7231 | 0.0118 | −0.185 |
| ■ BARCS + CNV | 0.9533 | 0.9762 | 0.7231 | 0.0118 | −0.042 |
| ■ Official MAGeCK-MLE | 0.9598 | 0.9795 | 0.7098 | 0.0074 | −0.207 |
| ■ Official MAGeCK-MLE + CNV | 0.9501 | 0.9762 | 0.7098 | 0.0074 | −0.006 |

The endpoint results in Table S5 show that CNV correction reduces the intended nuisance association, from 0.185 to 0.042 for BARCS and from 0.207 to 0.006 for MAGeCK-MLE, but decreases AUROC for both methods. After correction, the BARCS–MAGeCK AUROC difference is 0.00316 (paired stratified-bootstrap 95% CI 0.00265 to 0.00924), and the average-precision difference is effectively zero (95% CI 0.00349 to 0.00295). At nominal FDR 0.05, BARCS has 1.33 percentage points more recall but also a larger realized FDP within the gold-standard set (0.0118 versus 0.0074). These data do not establish a ranking or calibration advantage for either count model.

